# Chromosomal dynamics predicted by an elastic network model explains genome-wide accessibility and long-range couplings

**DOI:** 10.1101/081182

**Authors:** Natalie Sauerwald, She Zhang, Carl Kingsford, Ivet Bahar

**Affiliations:** Computational Biology Department, School of Computer Science, Carnegie Mellon University, 3064 Pittsburgh, PA 15213; Department of Computational and Systems Biology, School of Medicine, University of Pittsburgh, 3064 Pittsburgh, PA 15213

## Abstract

Understanding the three-dimensional (3D) architecture of the chromatin and its relation to gene expression and regulation is fundamental to understanding how the genome functions. Advances in Hi-C technology now permit us to have a glimpse into the 3D genome organization and identify topologically associated domains (TADs), but we still lack an understanding of the structural dynamics of chromosomes. The dynamic couplings between regions separated by large genomic distances (> 50 megabases) have yet to be characterized. We adapted a well-established protein-modeling framework, the Gaussian Network Model (GNM), to the task of modeling chromatin dynamics using Hi-C contact data. We show that the GNM can identify structural dynamics at multiple scales: it can quantify the fluctuations in the positions of gene loci, find large genomic compartments and smaller TADs that undergo en-bloc movements, and identify dynamically coupled distal regions along the chromosomes. We show that the predictions of the GNM correlate well with DNase-seq and ATAC-seq measurements on accessibility, the previously identified A and B compartments of chromatin structure, and pairs of interacting loci identified by ChIA-PET. We describe a method to use the GNM to identify novel cross-correlated distal domains (CCDDs) representing regions of long-range dynamic coupling and show that CCDDs are often associated with increased gene coexpression using a large-scale analysis of 212 expression experiments. Together, these results show that GNM provides a mathematically well-founded unified framework for assessing chromatin dynamics and the structural basis of genome-wide observations.

## Introduction

The spatial arrangement of chromosomes within the nucleus plays a crucial role in gene regulation, cell replication and mutations (1–5). Recent experimental methods such as Hi-C (6) derived from chromosome conformation capture (3C) (7) have made it possible to characterize the physical contacts between gene loci on a genome-wide scale. These studies revealed hierarchical levels of organization, from large (so called “A” and “B”) compartments corresponding to active and inactive chromatin respectively (6), to smaller compact regions called topologically associated domains (TADs) (8). Hi-C-measured spatial relationships have been related to chromosomal alterations in cancer (9) and TADs have been pointed out to contain clusters of genes that are co-regulated (10). Interactions between sequentially (but not necessarily spatially) distant genes along the DNA 1-dimensional (1D) structure, termed *long-range interactions*, have been implicated in gene regulation—for example, distal expression quantitative trait loci (eQTLs) tend to be much closer in 3D space (11) to their target genes than expected by chance.

Several computational methods have contributed to these and other characterizations of chromosomal architecture (8,12–18). However, chromosome structure is dynamic and complex, and its exact nature and influence on gene expression and regulation remain unclear. The scale, complexity, and noise inherent in the available data make it challenging to determine exact spatial relationships and underlying chromatin architecture and its structure-based dynamics. In particular, long-range spatial interactions have proven difficult to characterize with Hi-C data, and most computational analyses attempt to identify a static chromosomal architecture despite its known dynamic nature. There have also been efforts to mathematically characterize the dynamics of the genome separate from its structure, particularly through describing the emergence of cell types during development as bifurcations from a stable equilibrium (19).

Chromatin structure is often described in terms of TADs, whose identification is a 1D problem: it involves searching for sequentially contiguous groups of highly interconnected loci along the diagonal of the Hi-C matrix of intra-chromosomal contacts. Spatial couplings between sequentially distant genomic regions, on the other hand, represent a new dimension to search and the identification of such long-range couplings is a more challenging problem. Several methods have sought to identify long-range interactions from 3C-based data (13,20–23), but the scale of these interactions is still small compared to that of the full chromosome. Most methods detect interactions within 1-2 Mbp, or up to 10Mbp (24), so extending the span of predicted long-range couplings to the order of tens of millions of base pairs may yield further insights into regulatory actions. Such long-range correlations may originate from physical proximity in space, or other indirect effects similar to those in allosteric structures. Assessment of such long-range correlations is important for gaining a better understanding of the physical basis of gene expression and regulation.

We adopt here the Gaussian Network Model (GNM), a highly robust and widely tested framework developed for modeling the intrinsic dynamics of biomolecular systems (25–27), and we adapt it to the topology-based modeling of chromosomal dynamics. The only input GNM requires is a map of 3D contacts. Here, this information is provided by Hi-C data, which gives contact frequencies between genomic loci. The Hi-C matrix is used for constructing the Kirchhoff (or Laplacian) matrix **Γ** which uniquely defines the equilibrium dynamics of the network nodes (genomic loci) as well as their spatial cross-correlations. Notably, the use of Laplacian-based graph segmentation has been recently shown to help identify topological domains from Hi-C data (28,29). Our approach differs in the method of construction of **Γ**, the inclusion of the complete spectrum of motions, and the application to a broad range of observables. We show, and verify upon comparison with an array of experimental data and genome-wide statistical analyses, that the GNM provides a robust description of accessibility to the nuclear environment as well as co-expression patterns between gene-loci pairs separated by tens of megabases. The analysis is mathematically rigorous, efficient, and extensible, and may serve as an excellent framework for drawing inferences from Hi-C and other advanced genome-wide studies toward establishing the structural basis of regulation.

## Results and Discussion

### Extension of the Gaussian Network Model to Modeling Chromatin Dynamics

The GNM has proven to be a powerful tool for efficiently predicting the equilibrium dynamics of almost all proteins and their complexes/assemblies which can be accessed in the Protein Data Bank (PDB) (30,31), and has been incorporated into widely used molecular simulation tools such as CHARMM (32). It is particularly adept at predicting topology-dependent dynamics and identifying long-range correlations — the type of modeling that has been a challenge in chromatin 3D modeling studies. Hi-C matrices, in which each entry represents the frequency of contacts between pairs of genomic loci, can be interpreted as chromosomal contact maps similar to those **(Γ)** between residues used in the GNM representation of proteins.

There are several differences between the Hi-C and GNM **Γ** matrices. The first is the size: human chromosomes range from ~50 to 250 million base pairs. When binned at 5kb resolution this leads to 10,000 – 50,000 bins per chromosome. GNM provides a scalable framework, where the collective dynamics of supramolecular systems represented by 10^4^–10^5^ nodes (such as the ribosome or viruses) can be efficiently characterized. GNM may therefore be readily used for analyzing intrachromosomal contact maps at high resolution. The second is the precision of the data. Experimental methods for resolving biomolecular structures such as X-ray crystallography, NMR, and even cryo-electron microscopy yield structural data at a much higher resolution than current genome-wide studies. The Hi-C method is population-based (derived from hundreds of thousands to millions of cells) and noisy. However, the GNM results are usually robust to variations in the precision/resolution of the data on a local scale, and require only the overall contact topology rather than detailed spatial coordinates, which supports the utility of Hi-C data. Third, the chromatin is likely to be less ‘structured’ than the structures at the molecular level, and it is likely to sample an ensemble of conformations that may be cell or context-dependent. Single-cell Hi-C experiments have indicated cell-cell variability in chromosome structure on a global scale, though the domain organization at the megabase scale is largely conserved (33). Therefore, structure-based dynamic features may be assessed at best at a probabilistic level. With these approximations in mind, we now proceed to the extension of GNM to characterize chromosomal dynamics (see **Fig. 1**).

**Fig. 1.**
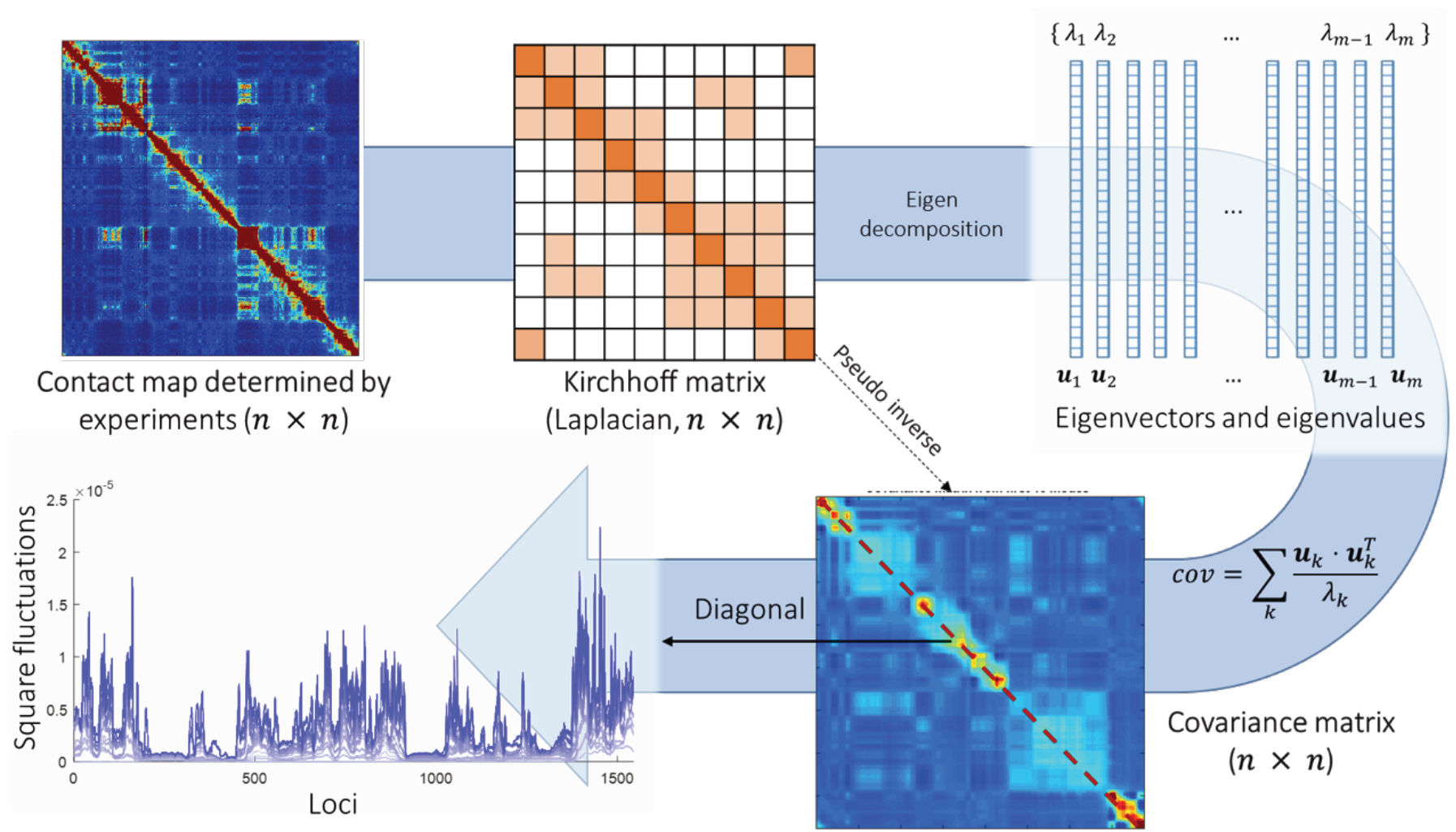
Schematic description of GNM methodology applied to Hi-C data. The interloci contact data represented by the Hi-C map (upper left, for *n* genomic bins (loci)) is used to construct the GNM Kirchhoff matrix, **Γ** (top, middle). Eigenvalue decomposition of **Γ** yields a series of eigenmodes which are used for computing the covariance matrix (lower, right), the diagonal elements of which reflect the mobility profile of the loci (bottom, left), and the off-diagonal parts provide information on locus-locus spatial cross-correlations. ***u***_*k*_, *k*th eigenvector; *λ*_*k*_, feth eigenvalue; *m,* number of nonzero modes, starting from the lowest-frequency mode, included in the GNM analysis (*m* ≤ *n* – 1). In the present application to the chromosomes, *n* varies in the range 10248 ≤ *n* **≤** 49850, the lower and upper limits corresponding respectively to chromosomes 22 and 1.

The GNM describes the structure as a network of beads/nodes connected by elastic springs. The network topology is defined by the Kirchhoff matrix **Γ**, whose elements are

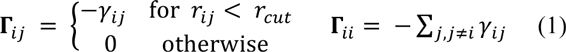

Here *γ*_*ij*_ represents the strength or stiffness of interaction between beads *i* and *j* (or the force constant associated with the spring that connects them), *γ*_*ij*_ is their separation in the 3D structure, and *r*_*cut*_ is the distance limit for making contacts (or for being connected by a spring). In the application to proteins, the beads represent the individual amino acids (*n* of them), their positions are identified with those of the *α*-carbons, and a uniform force-constant *γ*_*ij*_ = *γ* is adopted for all pairs (1 ≤ *i, j ≤ n*), with a cutoff distance of *r*_*cut*_ ~ 7Å. In the extension to human chromosomes, we redefine the network nodes and springs such that beads represent genomic loci consistent with the resolution of the Hi-C data. We set *γ*_*ij*_ equal to the Hi-C contact counts reported for the pair of genomic bins (15) *i* and *j* after normalization by vanilla coverage (VC) method (13) (see Methods and *Supplementary Information* SI). The elements **Γ**_*ij*_ is taken to be directly proportional to the actual number of physical nucleotide-nucleotide contacts between the loci *i* and *j*, which permits us to directly incorporate the strength of interactions in the network model. In a recent study, the Kirchhoff (Laplacian) matrix is normalized after construction (28,29), but we choose not to because it removes the information of packing density of nodes, renders the calculation of square fluctuations meaningless, and disables the comparison with chromatin accessibility.

The cross-correlation between the spatial displacements of loci *i* and *j* is obtained from the pseudoinverse of **Γ**, evaluated as

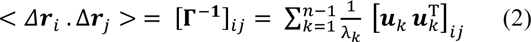

where the summation is performed over all modes of motion intrinsically accessible to the network, obtained by eigenvalue decomposition of **Γ**. The respective frequencies and shapes of these modes are given by the *n*-1 nonzero eigenvalues (*λ*_*k*_) and corresponding eigenvectors (***u***_*k*_). Cross-correlations are organized in the ***n×n*** covariance matrix, **C**. The diagonal elements of **C** are the predicted mean-square fluctuations (MSFs) in the positions of the loci under physiological conditions, also called the *mobility profile* of the chromosomes. The eigenvectors are *n*-dimensional vectors representing the normalized displacements of the *n* loci along each mode axis, and 1/*λ*_*k*_ rescales the amplitude of the motion along the *k*^*th*^ mode. Lower frequency modes (smaller *λ*_*k*_) make higher contributions to observed fluctuations and correlations; they usually embody large substructures if not the entire structure, hence their designation as *global modes*. This is in contrast to high frequency modes, which are highly localized, and often filtered out to better visualize cooperative events. See SI for more information.

### Loci dynamics correlate well with experimental measures of chromatin accessibility

We first evaluated the mobility profiles of the chromosomes for GM12878 cells from a human lympho-blastoid cell line with relatively normal karyotype. **Fig. 2** shows the MSFs obtained with the GNM (*blue curves*) for the loci on three chromosomes (1, 15 and 17, in respective panels ***A*, *B*** and ***C***). Results for all other chromosomes are presented in *Supplementary* **Fig. S1**.

**Figure 2.**
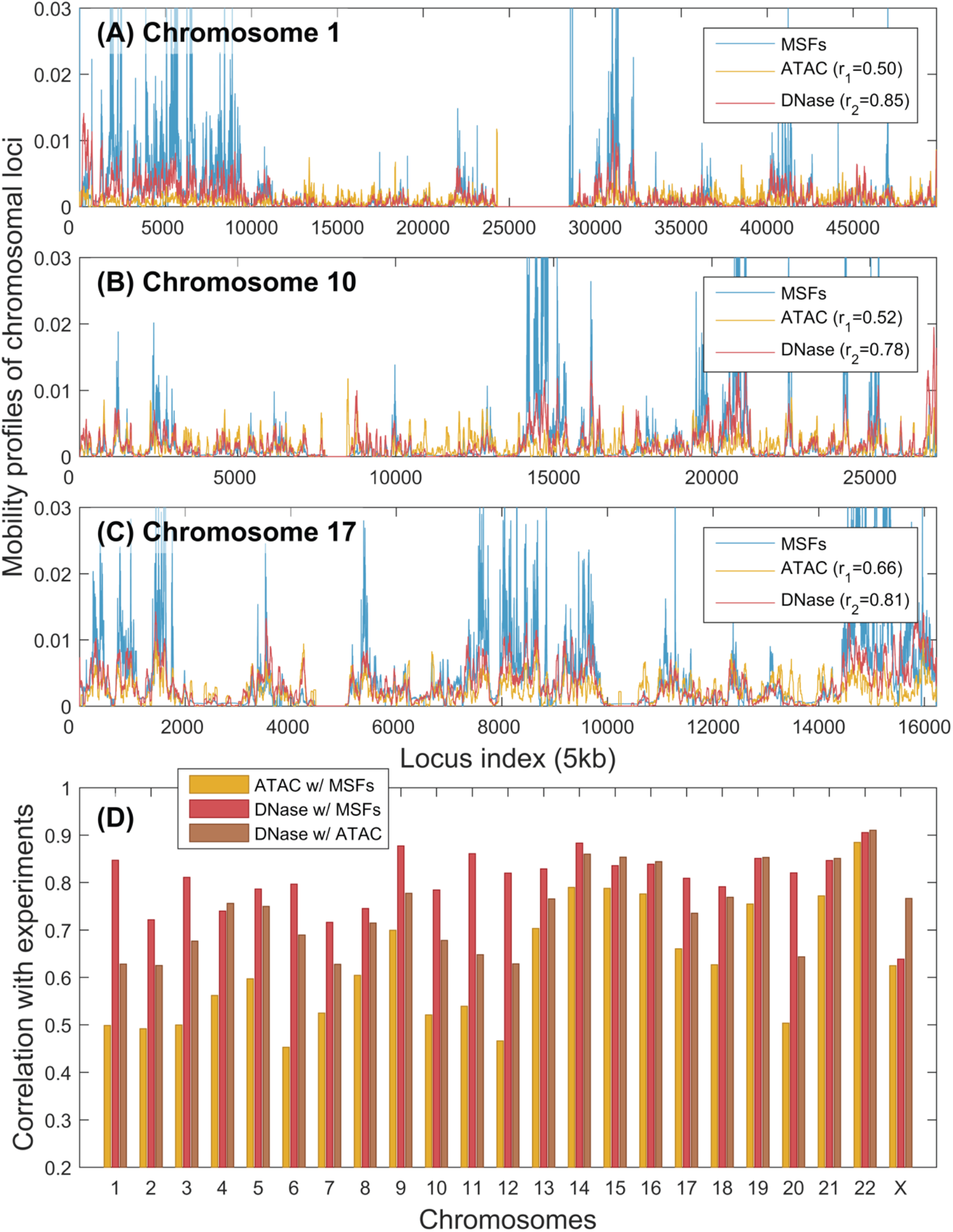
Correlations between GNM-predicted mobilities of chromosomal loci and data from chromatin accessibility experiments. (A) – (C) Mobility profiles (MSFs of loci) obtained from GNM analysis of the equilibrium dynamics of chromosomes 1, 17, and X, respectively, shown in blue, are compared to the DNA accessibilities probed by ATAC-seq (yellow) and DNA-seq (red) experiments. GNM results are based on 500 slowest modes. *r*_*1*_ is the Spearman correlations between GNM predictions and DNase-seq experiments; and *r*_*2*_ is that between GNM and ATAC-seq. (D) Spearman correlations between theory and experiments for all chromosomes (red and yellow bars, as labeled). The Spearman correlation between the computed MSFs and experimental ATAC-seq data averaged over all chromosomes is 0.62 ± 0.13, and that between MSFs and DNase-seq data is 0.81 ± 0.06. For comparison, we also display the Spearman correlation between the two sets of experimental data (brown bars); the average in this case is 0.74 ± 0.09.

GNM application to H/D exchange data has shown that the MSFs of network nodes can be directly related to the accessibility of the corresponding sites: exposed sites enjoy higher mobility, and those buried have suppressed mobilities (39). The entropic cost of exposure to the environment for a given site can be shown to be inversely proportional to its MSF based on simple thermodynamic arguments applied to macromolecules subject to Gaussian fluctuations (such as those represented by the GNM) (39). We examined whether GNM-predicted mobility profiles were also consistent with data from chromatin accessibility experiments. We compared our predictions with two measures of chromatin accessibility, DNase-seq (40) and ATAC-seq (35), shown respectively by the *yellow* and *red* curves in **Fig. 2 *A-C***.

**Fig. 2** shows that the MSFs of chromosomal loci, predicted by the GNM, are in very good agreement with the accessibility of loci as measured by DNase-seq. The corresponding Spearman correlations for the three chromosomes illustrated in panels ***A-C*** vary in the range 0.78-0.85 (see inset), and the computations for all 23 chromosomes (panel ***D***, yellow bars) yield an average Spearman correlation of 0.807 (standard deviation of 0.062). The average Spearman correlation between GNM MSFs and ATAC-seq data is somewhat lower: 0.623 ± 0.126. Interestingly, the average Spearman correlation between the two sets of experimental data was 0.741 ± 0.089, suggesting that the accuracy of computational predictions is comparable to that of experiments, and that the DNase-seq provides data more consistent with computational predictions. ATAC-seq maps not only the open chromatin, but also transcription factors and nucleosome occupancy (41), which may help explain the observed difference.

These results show that the mobility profiles predicted by the GNM for the 23 chromosomes accurately capture the accessibility of gene loci. The agreement with experimental data lends support to the applicability and utility of the GNM for making predictions on chromatin dynamics. The current results were obtained by using subsets of *m* = 500 GNM modes for each chromosome, which essentially yield the same profiles and the same level of agreement with experiments as those obtained with all modes (see *Supplementary* **Fig. S2**). The use of a subset of modes at the low frequency end of the spectrum improves the efficiency of computations, without compromising the accuracy of the results. Computations repeated for different levels of resolution (from 5kb to 50kb per bin) also showed that the results are insensitive to the level of coarse-graining (*Supplementary* **Fig. S3**) which further supports the robustness of GNM results.

### Domains identified by GNM at different resolutions correlate with known structural features

Compartments, first identified by Lieberman-Aiden et al. (6), are multi-megabase-sized regions in the genome that correspond to known genomic features such as gene presence, levels of gene expression, chromatin accessibility, and histone markers. Hi-C experiments have revealed two broad classes of compartments: “A” compartments generally associated with active chromatin, containing more genes, fewer repressive histone markers, and more highly expressed genes; and “B” compartments, for less accessible DNA, sparser genes, and higher occurrence of repressive histone marks. TADs (8) are finer resolution groupings of chromatin distinguished by denser self-interactions and associated with characteristic patterns of histone markers and CTCF binding sites near their boundaries. The multiscale nature of GNM spectral analysis allows hierarchical levels of organization to be identified computationally, and it is of interest to examine to what extent these two levels can be detected.

As presented above, the GNM low frequency modes reflect the global dynamics of the 3D structure, and increasingly more localized motions are represented by higher frequency modes. We identified domains from subsets of GNM modes that group regions of similar dynamics (see Methods). In order to verify whether these dynamical domains correspond to TADs at various resolutions, we used the TAD-finder Armatus (14), varying its *γ* parameter that controls resolution. We measure the agreement between GNM domains and TADs using the variation of information (VI) distance, which computes the agreement between two partitions, and where a lower value indicates greater agreement (42). For each choice *k* of number of modes, the *γ*_*k*_ that minimizes the VI distance between the GNM domains and the Armatus domains was selected. This resulted in a mean VI value for optimal parameters of 1.251, significantly lower than the VI distance of 1.946 obtained when the GNM domains were randomly re-ordered along the chromosome and compared back to the original TADs (empirical p-value < 0.01 for all chromosomes). **Fig. 3*A*** shows the comparison for each chromosome between the VI value for the optimally matched TAD boundaries with the GNM domains and the distribution of VI values from the randomly shuffled domains. As the number of included GNM modes is increased, *γ*_*k*_ monotonically increases as well, showing that the number of GNM modes is a good proxy for the scale of chromatin structures sought (**Fig. S4**).

**Figure 3.**
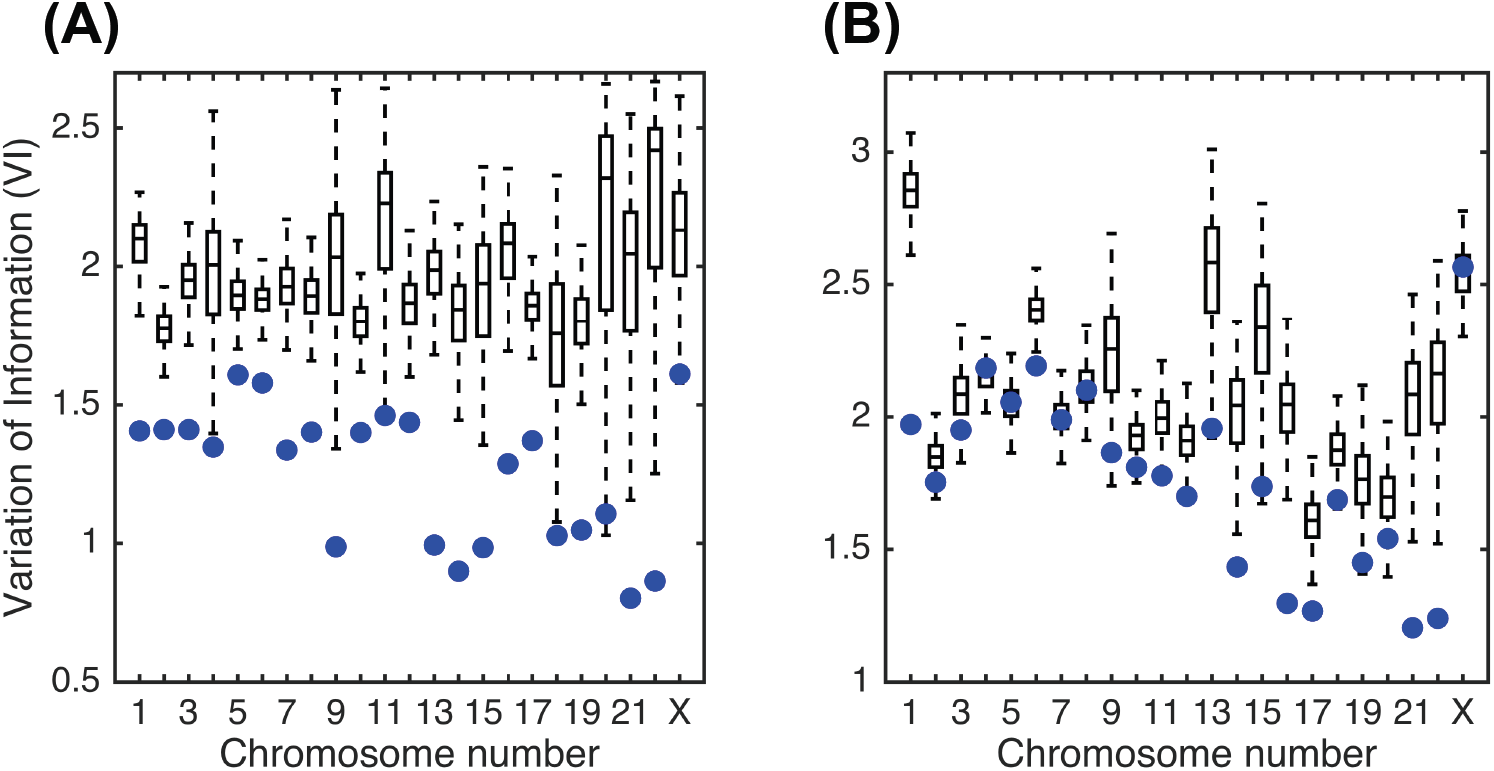
Variation of information (VI) measures for comparing GNM domains with (A) TADs and (B) compartments (lower VI indicates greater agreement). Box plots show the distribution of VI values obtained by randomly shuffling GNM domains and comparing to original TAD and compartment boundaries. Blue dots represent the VI value of the true GNM domains with TADs and compartments, respectively.

Furthermore, GNM predicts large-scale global motions using a relatively low number of modes, so we compared these to larger-scale compartments. We found that the first 5-20 non-zero modes correspond fairly well to compartments. For each chromosome, we selected the number of modes that produced the smallest VI distance between Lieberman-Aiden compartments and GNM domains. This yielded a mean optimal VI distance of 1.771 (using an average of 13 modes). This is significantly lower than the mean optimal VI distance of 2.088 when the locations of Lieberman-Aiden compartments are randomly shuffled along the chromosome, though the difference is only statistically significant for 16 of the 23 chromosomes, with p equal to 0.05. The comparisons of GNM domains with compartments for each chromosome can be seen in Fig. 3*B*. **Fig. S5** shows an example of the GNM domains found using the number of modes that minimizes the VI with compartments or TADs. The ability of GNM to recapitulate both TADs and compartments—two organizational levels of wildly different scales—shows the flexibility and generality of the GNM approach. We note that a TAD-finding method using only the second eigenpair (Fiedler value/vector) of the Laplacian has also been developed (28) and tested on 100kb resolution data. By including more eigenvectors, we are able to identify TADs closer to Armatus on all chromosomes (as measured by lower VI) at 5kb and for 18/23 chromosomes at 100kb resolution (see Figure **S6A** and **C**). Though the Fiedler vector-based method identifies compartments better at low resolution, their method performs poorly at finer resolution, while GNM remains robust to resolution changes. We are also able to identify compartment sets with lower VI on all chromosomes at 5kb (**Figure S6B** and **D**). Further corroborating the benefit of using multiple modes, it has been shown in early studies that spectral clustering by using more eigenvectors can outperform partitioning methods which only use one eigenvector (43,44).

### Loci pairs separated by similar 1D distance exhibit differential levels of dynamic coupling, consistent with ChIA-PET data

**Fig. 4** displays the covariance map generated for the coupled movements of the loci on chromosome 17 (of GM12878 cells), based on Hi-C data at 5 kb resolution. Panel *A* displays the cross-correlations (see equation 2) between all loci-pairs as a heat map. Diagonal elements are the MSFs (presented in **Fig. 2*C***). The curve along the upper x-axis in **Fig. 4*A*** shows the average cross-correlation of each locus with respect to all others; the peaks indicate the regions tightly coupled to all others, probably occupying central positions in the 3D architecture. Results for other chromosomes are presented in *Supplementary* **Fig. S7**. The covariance map is highly robust and insensitive to the resolution of the Hi-C data. The results in **Fig. 4*A*** were obtained using all the *m* = 15,218 nonzero modes corresponding to 5kb resolution representation of chromosome 17. Calculations repeated with lower resolution data (50kb) and fewer modes (500 modes) yielded covariance maps that maintained the same features (**Fig. S8**).

**Figure 4.**
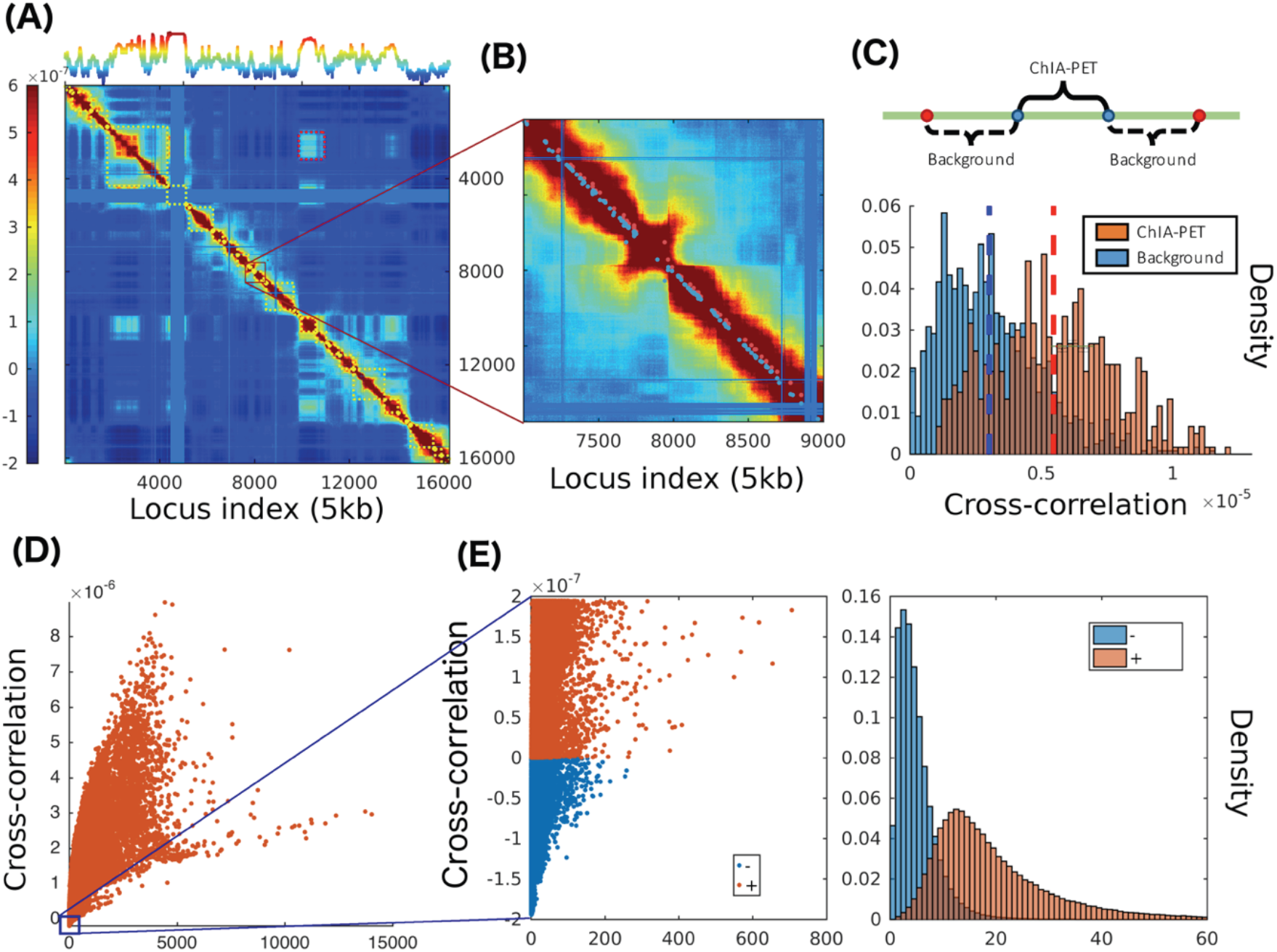
Covariance map computed for chromosome 17 and comparison with ChIA-PET data and contacts from Hi-C experiments. (A) Covariance matrix computed for chromosome 17, color-coded by the strength and type of cross-correlation between loci pairs ranged from 5th to 95th percentile of all cross-correlation values (see the color bar on the left). The curve on the upper abscissa shows the average overall offdiagonal elements in each column, which provides a metric of the coupling of individual loci to all others. The blocks along the diagonal indicate loci clusters of different sizes that form strongly coupled clusters. The red dashed boxes indicate the pairs of regions exhibiting weak correlations despite genomic distances of several megabases. The blue bands correspond to the centromere, where there are no mapped interactions. (B) Close-up view of a region along the diagonal. *Red dots* near the diagonal indicate pairs (separated by ~100 kb) identified by ChIA-PET to interact with each other; nearby *blue points* are control/background pairs. (C) Stronger cross-correlations of ChIA-PET pairs compared to the background pairs. (D) Dependence of cross-correlations on the number of contacts observed in Hi-C experiments. A broad distribution is observed, indicating the effect of the overall network topology (beyond local contacts) on the observed cross-correlations. (E) Loci pairs exhibiting anti-correlated (same direction, opposite sense) movements usually have fewer contacts, compared to those exhibiting correlated (same direction, same sense) pairs of the same strength.

Owing to their genomic sequence proximity, the entries near the main diagonal of the covariance map tend to show relatively high covariance values (colored *yellow-to-brown;* **Fig. 4*A***). Note that even the close vicinity of the diagonals (e.g. loci intervals of ≥ 200) represents (at 5 kb resolution) genomic loci separated by more than 1 megabase. The covariance map clearly shows that there are strong couplings between loci separated by a few megabases. We show an example of such regions in **Fig. 4*B***. While the loci pairs located in the *dark red band* along the diagonal appear all to exhibit strong couplings, a closer examination reveals differential levels of cross-correlations that are in good agreement with the data from Chromatin Interaction Analysis by Paired-End Tag Sequencing (ChIA-PET) experiments (45). The ‘long-range’ interactions identified by ChIA-PET (36) are indicated in panel ***B*** by *red dots* (close to the diagonal). These are interacting loci separated by several hundreds of kb. We selected background pairs separated by the same 1D distance, on both sides of the ChIA-PET pair, and compared the cross-correlations predicted for the two sets along each chromosome (**Fig. 4*C***). The background pairs (*blue bars*) show weaker GNM cross-correlations compared to the ChIA-PET pairs (*red bars*) although they are separated by the same genomic distance along the chromosome.

Similar statistical analysis repeated for all 23 chromosomes showed that the cross-correlations of ChIA-PET pairs were greater than those of background pairs of the same genomic distance on every chromosome, with all p-values less than 10^−19^.

### Cross-correlations between loci motions are global properties that result from the overall chromosomal network topology

In general, loci-loci cross-correlations become weaker with increasing distance along the chromosome, and some pairs show anticorrelations (i.e. move in opposite directions; see scale bar in **Fig. 4*A***). Yet, we can distinguish distal regions that exhibit notable cross-correlations in the spatial movements (off-diagonal lighter-colored blocks). The levels of cross-correlations do not necessarily need to scale with the interaction strengths between the correlated loci (or number of contacts detected by Hi-C). On the contrary, a broad range of cross-correlations is observed for a given number of contacts, indicating that the observed correlations are global properties defined by the entire network topology and reflect the collective behavior of the entire structure. **Fig. 4*D*** displays the computed cross-correlations as a function the number of contacts, showing that some pairs of loci display much stronger correlations revealed by the GNM than others that make more Hi-C contacts. **Fig. 4*E*** shows that the anticorrelated pairs of loci (*blue*) usually have fewer contacts than those (*red*) exhibiting positive cross-correlations of the same strength.

### Distal regions predicted to be strongly correlated in their spatial dynamics exhibit higher co-expression

The GNM covariance map further shows correlations between farther apart (>10 Mbp) regions. In contrast to the main diagonal, the majority of the off-diagonal space typically shows significantly weaker correlations. Regions in this space with higher than expected covariance values represent dynamically linked windows along the chromosome, which may represent long-range interactions. We call these pairs of windows *cross-correlated distal domains* (CCDDs). To identify CCDDs, we set a threshold for each covariance matrix equal to the absolute value of the minimum covariance. Treating the remaining adjacent pairs as edges in a graph, we then locate connected components beyond the widest section of the main diagonal and above the threshold that contain more than one bin pair, and find the maximal-area rectangle contained within each connected region of high covariance values (see **Fig. S9**). These CCDDs are therefore pairs of regions distant along the chromosome, composed each of highly interconnected loci, which also exhibit relatively high cross-correlations compared to other regions of similar genomic separation. Previous methods for identifying long-range chromatin interactions (13,20,21,45) have focused on locating individual points of interaction within 1-2 Mbp apart, while CCDDs tend to be on the order of tens of Mbp apart and supported by groups of interacting loci.

Highly distant gene pairs within CCDDs show greater co-expression values than gene pairs outside these regions (p-value < 10^−7^ using the background defined below). For each CCDD, we identified the genes contained within the region and measured the co-expression of each gene pair from distant chromosomal segments. The background gene pairs were gathered from outside the CCDDs but with similar genomic separation as the CCDD gene pairs. We computed gene expression correlations from 212 experiments (see Methods and *Supplementary* Table S1).

As seen in **Fig. 5**, the CCDDs containing specifically gene pairs that are between 50 and 100 Mbp apart are much more highly co-expressed than background gene pairs at the same genomic distance (p-value < 10^−19^). This indicates that the dynamic coupling of these genes, as revealed by GNM, may often be biologically important. CCDDs at smaller genomic distance (< 50 Mbp) exhibit similar co-expression distributions to the background gene pairs, likely due to the effect of shorter genomic distances including more co-regulated genes within the background. Beyond distances of 100 Mbp, there are not sufficient gene pairs within CCDDs to draw any meaningful conclusions. Dynamically coupled regions that are very distant sequentially but biologically linked through gene expression are therefore identifiable using the GNM covariance matrix.

**Figure 5.**
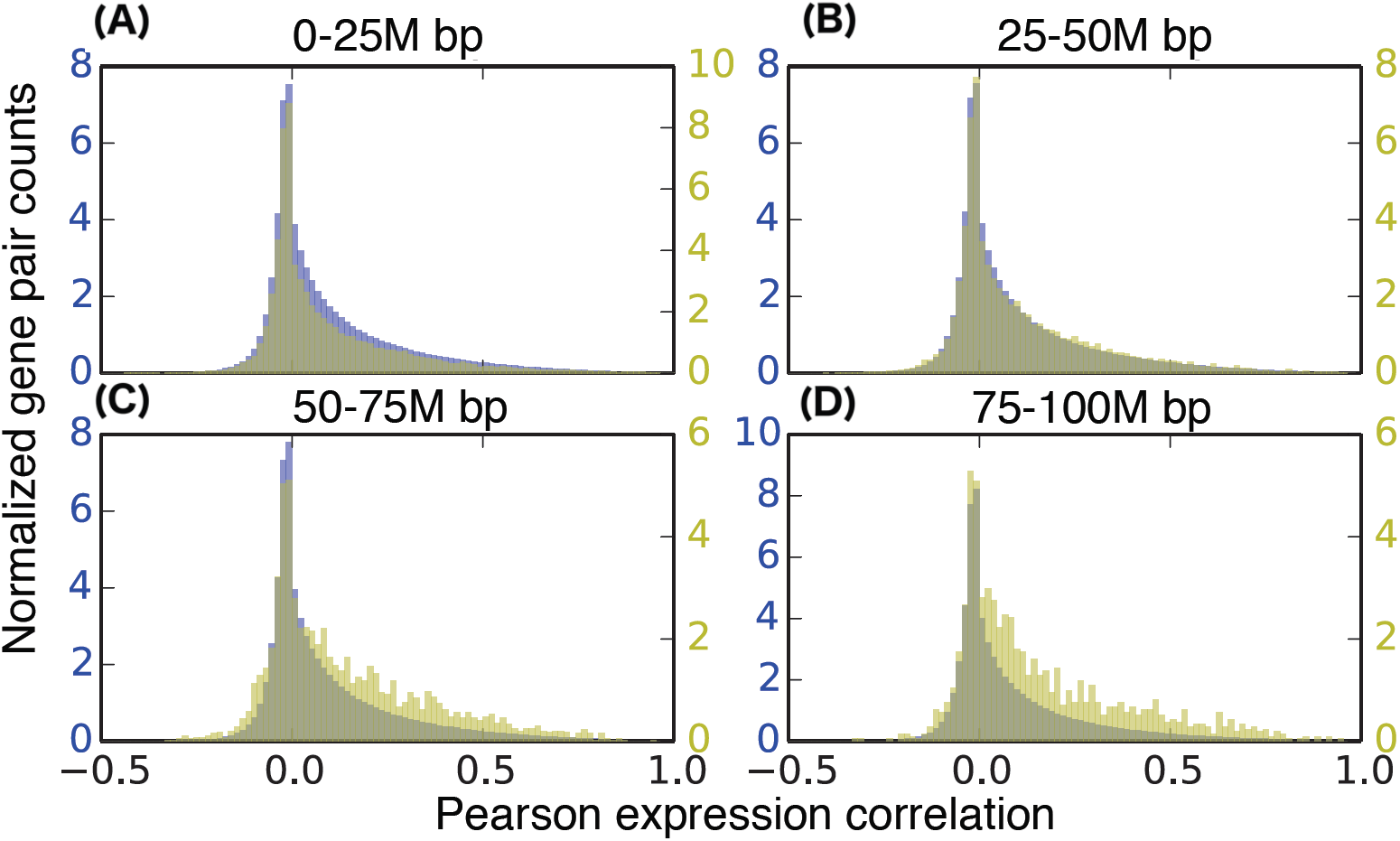
Correlating gene co-expression with CCDDs. In each histogram, the yellow distribution represents gene pairs from CCDDs and the blue distribution represents background gene pairs. All are showing the normalized number of gene pairs with a particular Pearson expression correlation for gene pairs within a distance of (A) 0-25 million base pairs, (B) 25-50 million base pairs, (C) 50-75 million base pairs, and (D) 75100 million base pairs. The more distant pairs (50-100 million base pairs apart) within the CCDDs show enriched expression correlations as compared to the background pairs. There were not enough gene pairs within CCDDs more than 100M base pairs apart to draw significant conclusions.

## Conclusions

This work represents the first analysis of chromosome dynamics using an elastic network model, GNM, which has found wide applications in molecular structural biology. Though other models (28,29) have examined genome structure through graph theoretical methods, we show that inclusion of the complete spectrum of motions in the analysis provides a more realistic picture of chromosomal dynamics in accord with a wealth of experimental data. The model proves capable of capturing various properties of chromosomes and permits the study of previously unidentified, highly distant regions with enriched co-expression. Individual displacements of 5kb resolution chromosome segments predicted by the GNM MSFs correlate very well with measures of chromatin accessibility. Furthermore, regulatory interactions discovered by ChIA-PET sequencing are explained by the strong cross-correlation values predicted by the GNM, and dynamic domains of various sizes deduced from GNM correspond to both compartments and TADs, two well-known structural elements of chromatin. This unifying framework further led to the discovery of biologically significant, dynamically coupled regions, termed CCDDs, which are sequentially extremely distant.

In general, the evaluation of dynamic features using structure-based models becomes prohibitively expensive with increasing size of the structure, hence the development of coarse-grained models and methods for exploring supramolecular systems dynamics. The chromatin size is well beyond the range that can be tackled efficiently by structure-based methods and realistic force fields. The applicability of the GNM to modeling chromatin dynamics lies in its ability to solve for the collective fluctuations and cross-correlations based on network contact topology, exclusively. No knowledge of structural coordinates is needed, nor do we predict structural models – a task that has been undertaken successfully by recent studies (17,18,46–54). We characterize the collective dynamics encoded by the overall chromosomal contact topology, driven by entropy, consistent with the ensemble-based properties of the genome structure. The method is extremely efficient. For example, GNM averages a real computing time of 1.5 hours per chromosome at 5kb resolution using 10 CPUs, and no multiple runs are needed, since a unique analytical solution is obtained for each system. The computing time is further shortened when lower resolution data are used: all GNM computations are performed within one minute for every chromosome at the resolution of 50kb. The efficiency of the computations permits a systematic study of different types of cell lines as well as the extension of the methodology to the entire set of interchromosomal contacts, rather than individual chromosomes.

Future GNM analyses of chromatin dynamics could focus on the nature of the long-range couplings, analysis of their biological significance, or the meaning of genomic regions that exhibit high covariances. GNM also predicts a measure of overall coupling of each genomic locus to others (see the curve along the upper x-axis in **Fig. 4*A***), the significance of which requires further investigation. The GNM was shown to capture several biological properties of chromosomes, but further insights on cooperative events, including the interchromosomal (*trans*) interactions is within reach by focusing on the softest (lowest frequency) modes of motion predicted by the GNM. Finally, advances in 3D embeddings of Hi-C data may open the way to adopting the Anisotropic Network Model (ANM) (55–57) for efficient modeling and visualization of the whole chromatin dynamics.

The present study is performed on GM12878 cells, but the GNM can be readily used for analyzing different cell types provided that Hi-C data are available, and the comparative analysis of the fluctuation spectrum and CCDDs can reveal the differences across cell types. Our preliminary analysis of the equilibrium fluctuations of chromosome 17 for five other cell types (K562, KBM7, IMR90, HUVEC, and NHEK) indeed showed similarities between the MSFs of gene loci as well as their cross-correlations, although some notable differences were also seen, e.g. the mobility profile for the epidermal cell line, NHEK, exhibited distinctive patterns at selected regions. Further work is needed to understand the biological significance of the observed heterogeneities in the genome-wide fluctuation spectrum of the different types of cells. Examination of structural variabilities across orthologous proteins and their mutants revealed close similarities between evolutionary changes in structure and the intrinsic dynamics of proteins (58). Conversely, ANM-predicted global dynamics conforms to the principal changes in structure across different forms of the same protein (59,60), and thus explains the structural adaptability of the protein to different functional states (61). It would be of interest to explore whether cell-cell variabilities as well differences in disease *vs* normal states could equally be rationalized in the light of chromatin dynamics as more data become accessible on cell-specific 3D genome organization.

## Methods

### Datasets

Our Hi-C data came from the large, high-resolution Hi-C dataset (13) (GEO accession GSE63525), pre-processed using vanilla coverage (VC) normalization (13) (See SI for comparison of normalization methods). We used 5 kb resolution, the highest available in this dataset for GM12878 cells, unless otherwise noted. The DNase-seq data were also from GM12878 cells collected as part of the ENCODE project (ENCFF000SKV) (34). The ATAC-seq measurements (35) were collected also on GM12878 cells (GEO accession GSM1155959). For both of these experimental datasets, bed-formatted peak files were downloaded from the study authors and the data was binned to the same resolution as the Hi-C data by adding all peak values within each bin. The binned data was then smoothed using moving average with a window size of 200kb. The long-range interactions from ChIA-PET were from ENCODE (ENCFF002EMO) (36). We used a two-sample T-test assuming unequal variances to quantify the difference between the covariance distributions of ChIA-PET and background interactions.

### GNM Domain Identification

For any set of modes, described by the corresponding set of eigenpairs, GNM domain boundaries are located by the sign changes of each of the eigenvectors. These eigenvectors are often noisy, so we first smooth them with local regression using weighted linear least squares and a first-degree polynomial model. The smoothing window was chosen to be the smallest value that minimizes the number of domains of length one, where a domain of length one is defined as a domain that begins and ends in the same bin. The sign changes of the smoothed eigenvectors represent changes in directionality of motion and are labeled hinge sites. Each GNM domain is therefore a region between the union of hinge sites from each mode.

### Co-expression calculation

In order to calculate co-expression values for genes in this cell type, we downloaded every publically available RNA-seq experiment on GM12878 cells from the Sequence Read Archive (37), which gave 212 data sets. This raw read data was quantified using Salmon (38), resulting in 212 transcripts per kilobase million (TPM) values for every gene. Co-expression was then measured as the Pearson correlation of the two vectors of TPM values for a given gene pair.

## Acknowledgements

We thank Dr. C. S. Chennubhotla for the useful discussion with him. This research is funded in part by the NIH grants R01 GM099738 and U54 HG008540 to IB; and the Gordon and Betty Moore Foundation’s Data-Driven Discovery Initiative through Grant GBMF4554, National Science Foundation (CCF-1256087, CCF-1319998) and NIH (R01 HG007104) grants to CK. CK received support as an Alfred P. Sloan Research Fellow.

